# Lymphatic Endothelial Cells Regulate Neutrophil Phenotypes and Function in a Microphysiological Model of Infection

**DOI:** 10.64898/2026.03.24.714048

**Authors:** Kelsea A Sholty, Sheena C Kerr, David J Beebe

## Abstract

Early skin inflammation requires coordinated immune regulation, with neutrophils acting as first-line responders. While the blood vasculature and its role in neutrophil recruitment during infection has been extensively studied, the lymphatic system remains comparatively understudied despite its known role in immune cell trafficking. Growing evidence suggests lymphatic vessels actively participate in regulating inflammatory responses, yet whether they coordinate neutrophil behavior during skin infection remains unclear. *Staphylococcus aureus* is particularly problematic in this context, employing multiple immune evasion strategies and representing a major driver of antibiotic-resistant skin and soft tissue infections worldwide. To address this gap, we developed a human-based 3D microphysiological system incorporating luminal lymphatic endothelial vessels, a collagen matrix and bacteria to model an infected microenvironment. We evaluated neutrophil migration, phagocytosis and NETosis in response to *Escherichia coli* and *S. aureus*. Lymphatic endothelium amplified neutrophil migration in a bacterial-dependent manner, with *E. coli* promoting directional migration toward the vessel while *S. aureus* suppressed migration and directionality despite increased phagocytic uptake. *S. aureus* also induced myeloperoxidase-positive NETs with nuclear morphology consistent with vital NETosis, rescued by DNase treatment. To our knowledge, this is the first demonstration that lymphatic endothelium directly drives neutrophil behavior during skin infection.

## Introduction

Skin, the body’s largest organ, undergoes a tightly regulated healing process following injury, consisting of inflammation, regeneration and maturation. This process can be disrupted by microbial overgrowth, often resulting in infection and impaired healing. The growing prevalence of antibiotic-resistant infections poses a serious threat to public health and if not effectively controlled, these infections can escalate to sepsis and death^1,2^. A hallmark of infection-related complications is unregulated inflammation, which can further damage tissues and hinder repair^3,4^. Neutrophils are among the first immune responders to tissue injury and, under normal conditions, play a key role in controlling infection and promoting repair^5,6^. Some pathogens, namely *Staphylococcus aureus*, have evolved strategies to evade neutrophil killing, enhancing their ability to spread, making it one of the most prevalent bacteria in skin and soft tissue infections^7–11^. Notably, *S. aureus* is responsible for a significant proportion of antibiotic-resistant infections, contributing to considerable morbidity and mortality globally^1,12^. In addition to phagocytosis and degranulation, neutrophils respond to infection by releasing neutrophil extracellular traps (NETs). NET formation can occur through distinct processes broadly categorized as suicidal NETosis, which is associated with neutrophil cell death, and vital NETosis, in which neutrophils remain viable following chromatin release. While NET formation can aid microbial containment, excessive or poorly regulated NETosis has been linked to tissue damage and chronic inflammation, particularly in *S. aureus* infections^10,11^. Effective resolution of these neutrophil-driven inflammatory responses depends on coordinated immune regulation, a process in which the lymphatic vasculature is thought to play a central role.

The lymphatic system is thought to regulate neutrophil clearance and coordinate immune responses, offering a potential mechanism for controlling excessive inflammation and infection^13,14^. In other systems like the gut, the lymphatic system is known for its regulatory role in microbiome stability through the coordination of immune responses^15,16^. This emphasizes an important and largely unexplored area in understanding whether the lymphatic system plays a similar regulatory role in the skin, particularly following injury. In parallel, the skin microbiome is increasingly recognized for its role in modulating immune responses and presents a promising therapeutic avenue to restore balance and promote healing^17,18^. To study these interactions, we developed a physiologically relevant model using the LumeNEXT platform. This 3D, human microphysiological system (MPS), enables the formation of organized, lumen structures within an extracellular matrix (also referred to as biomimetic vessels). Unlike 2D cultures, LumeNEXT better mimics native tissue architecture and facilitates dynamic interactions between immune cells and microenvironmental cues. The platform has been successfully used to study immune cell interactions with both blood and lymphatic vessels, as well as cancer systems, demonstrating improved physiological relevance and versatility. Notably, interaction with a blood endothelial lumen has been shown to extend neutrophil lifespan to approximately 16–24 hours^19–23^. In this study, the MPS was designed to incorporate key elements of an infected microenvironment, including a lymphatic luminal endothelium, neutrophils and bacteria. Using this system, we observed that lymphatic endothelium enhanced neutrophil migration during bacterial challenge, with *E. coli* promoting strong directional migration while *S. aureus* suppressed migration and directionality supporting its greater virulence.

## Results

### Role of lymphatic system on neutrophil migration in MPS wound microenvironment

Neutrophils respond rapidly to bacterial infection, however the role of the lymphatic endothelium in this process is not well defined. To investigate lymphatic-neutrophil interactions, we developed an MPS that recapitulates a lymphatic endothelial vessel. The MPS is based on the LumeNEXT platform^24^, with PDMS device layers placed above and below two PDMS rods. When the central chamber is filled with type I collagen and polymerized, removal of the rods creates molded lumens that can be seeded with lymphatic endothelial cells to form a vessel or left unseeded to permit the addition of neutrophils into the system. The lymphatic MPS was previously characterized by demonstrating expression of lymphatic markers, barrier properties characteristic of lymphatic endothelium and secretion of growth factors and cytokines^20^. Here, we have adapted this model to incorporate a bacterial stimulus to permit study of neutrophil migration in bacterial infection (Fig. 1). pHrodo *E. coli* bioparticles were embedded into the collagen matrix as the bacterial component. These particles fluoresce green when phagocytosed permitting quantification of phagocytosis by neutrophils. *E. coli* was used as a representative bacterial stimulus that elicits neutrophil responses without the strong immune evasion strategies associated with *S. aureus*. One MPS lumen was seeded with primary human dermal lymphatic endothelial cells (HdLECs) (Fig. 2). To assess neutrophil migration and phagocytic activity, 32,000 neutrophils were added to the empty lumen and allowed to migrate for 4 hours prior to imaging. For neutrophil nuclear size analyses, devices were imaged at 24 hours. Single-cell quantification was performed using StarDist, a deep learning-based nuclear segmentation tool. Representative images show the device in the absence (Fig. 2A) and presence (Fig. 2B) of a lymphatic vessel, with bacteria present in both conditions. Results showed that neutrophil migration increased significantly when both bacterial components and lymphatic vessel were present compared to controls lacking either bacteria (*P* < 0.0001) or lymphatic vessel (*P* = 0.0128), with similarly low migration observed in both control conditions (Fig. 2C). Neutrophils showed a clear preference for migrating toward the lymphatic vessel in the presence of bacteria (Fig. 2B), with significantly more neutrophils migrating towards the lymphatic vessel (*P* < 0.0001) (Fig. 2D). Quantification of distance migrated demonstrated that neutrophils travelled further toward the lymphatic vessel compared to away from it when both bacteria and the vessel were present (Fig. 2E; representative donor shown, *P* = 0.0056), a finding consistent across donors (Supp. Fig. 1). This preferential migration was lost when bacteria were omitted from the system, suggesting a coordinated response mechanism between neutrophils and lymphatic endothelial cells within an infected microenvironment (Supp. Fig. 1).

**Figure 1.**
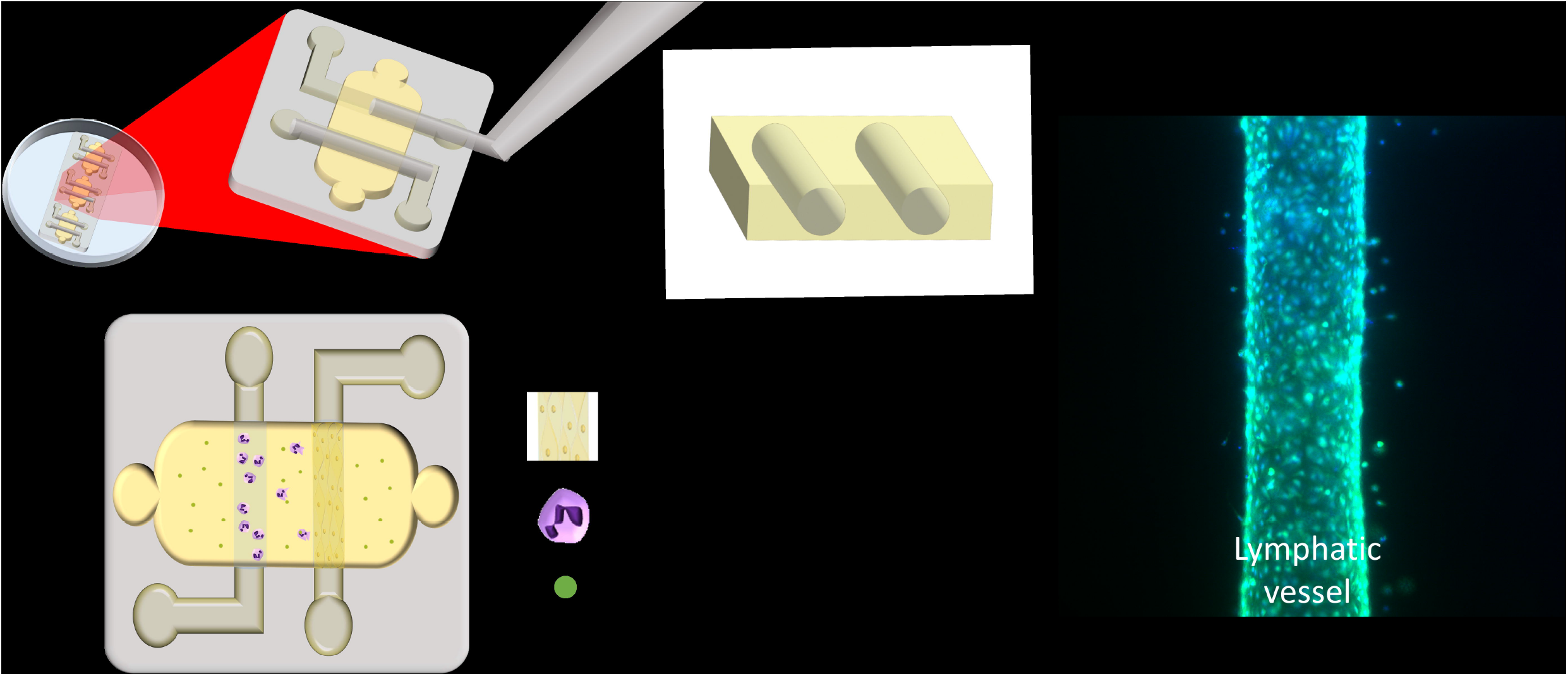
Microphysiological system design for modeling lymphatic-neutrophil interactions. (A) PDMS devices fabricated from soft lithography molds assembled onto MatTek glass-bottom culture dishes with integrated removable PDMS rods, and resultant lumen channels following rod removal after collagen polymerization. (B) Schematic showing the lumen seeding arrangement with one lumen seeded with human dermal lymphatic endothelial cells (HdLECs, yellow) to form a biomimetic lymphatic vessel and the adjacent lumen loaded with neutrophils (purple). pHrodo-conjugated bacterial bioparticles (green) are embedded within the surrounding collagen matrix. (C) Representative fluorescence image of a biomimetic lymphatic vessel formed within the collagen matrix.

**Figure 2.**
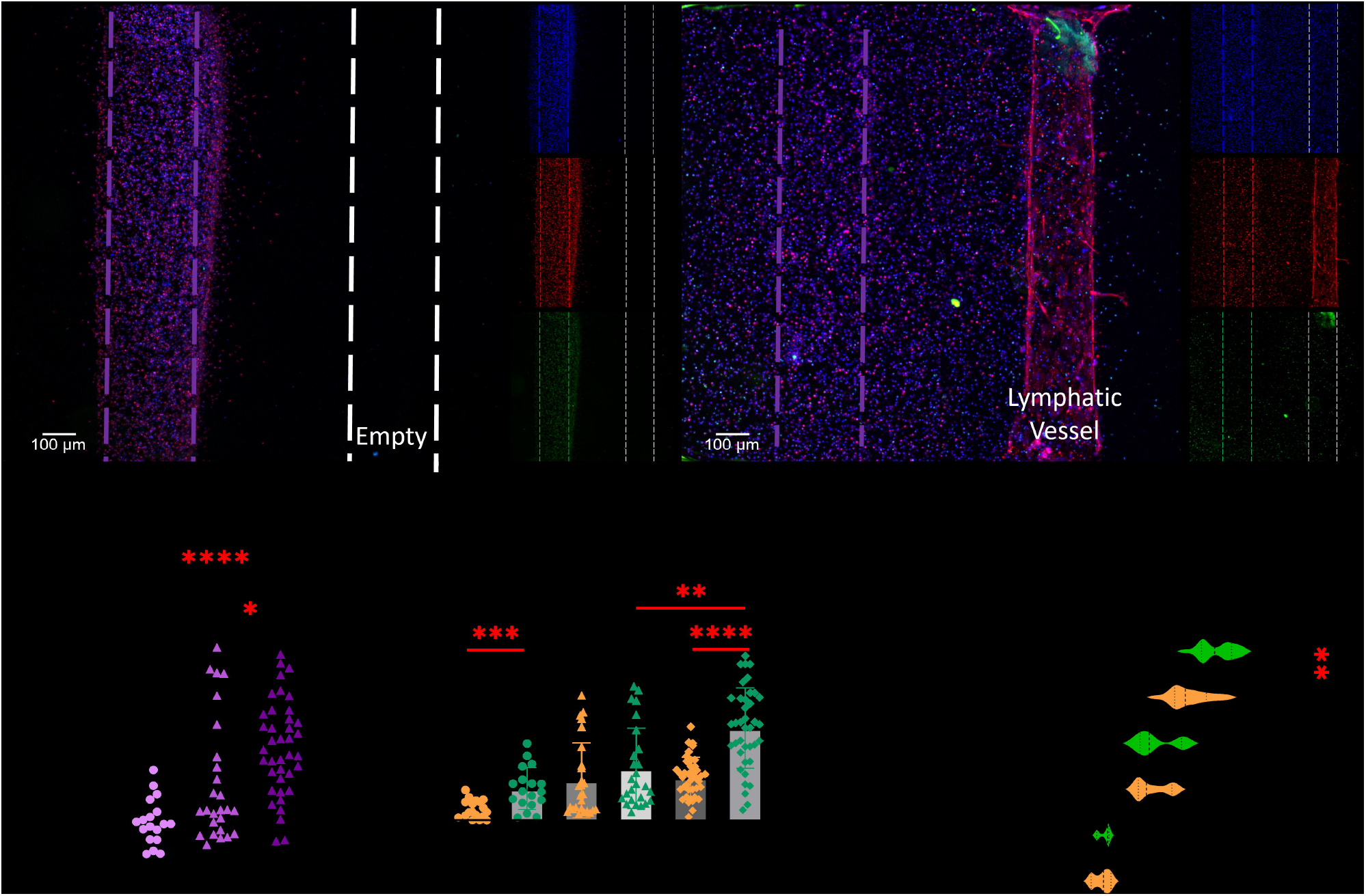
Lymphatic endothelial cells amplify neutrophil migration in a 3D infection model. (A) Representative fluorescence image showing minimal neutrophil migration (DAPI, blue) in the presence of *E. coli* pHrodo bioparticles (green) without a lymphatic vessel. (B) Representative fluorescence image showing increased neutrophil migration toward the lymphatic vessel (Phalloidin, red) in the presence of *E. coli* pHrodo bioparticles. Blue – DAPI; Red – Phalloidin Texas Red; Green – pHrodo (bacterial uptake). (C) Total neutrophil counts within the collagen matrix at 4 hours across three conditions, demonstrating that migration is significantly increased only when both bacteria and a lymphatic vessel are present. (D) Proximal (P) versus distal (D) neutrophil counts relative to the lymphatic vessel lumen at 4 hours, showing preferential migration toward the vessel in the presence of bacteria. (E) Representative donor migration distance across conditions at 4 hours, confirming greater distance traveled toward the lymphatic vessel when both components are present. All experiments were performed using neutrophils isolated from n = 4 independent donors. Data in C and D represent mean ± SD; data in E are displayed as violin plots. Between-condition comparisons performed using Kruskal-Wallis with Dunn’s post-hoc test; proximal versus distal comparisons performed using Mann-Whitney test. *p < 0.05, **p < 0.01, ***p < 0.001, ****p < 0.0001; ns, not significant.

### Neutrophil migration patterns between different bacteria

Neutrophil responses are coordinated and shaped by tissue context and microbial cues^5,6,25^. In skin, lymphatic endothelial signaling further modulates these behaviors^26–28^, though such interactions are rarely captured in real time. To evaluate how different bacterial species influence neutrophil behavior under these conditions, we assessed neutrophil migration, phagocytosis and nuclear morphology in response to either *Escherichia coli* or *Staphylococcus aureus*, the most common cause of skin and soft tissue infections. MPS were built as described above containing a biomimetic lymphatic vessel embedded within a collagen matrix. Within the collagen matrix, pHrodo-conjugated *E. coli* or *S. aureus* bioparticles served as a bacterial stimulus and enabled quantification of neutrophil phagocytosis visualized by green fluorescence following uptake. 32,000 neutrophils were added into the adjacent empty lumen and incubated for 4 hours before analysis using fluorescence microscopy. Using StarDist, a deep learning-based approach for automated single-cell nuclear segmentation in fluorescent images, we obtained single-cell measurements of neutrophil migration, fluorescence intensity and nuclear size (Fig. 3). Results revealed that significantly fewer neutrophils migrated in the presence of *S. aureus* compared to *E. coli* (*P* = 0.0019), with migration under *S. aureus* conditions indistinguishable from the no-bacteria control (*P* = 0.8646). (Fig. 3A). When considering directionality, preferential migration toward the lymphatic vessel was significant in the presence of *E. coli* (*P* = 0.0003) but was not observed under *S. aureus* conditions (*P* = 0.0983) (Fig. 3B). Neutrophils exposed to *S. aureus* displayed larger nuclear size than those responding to *E. coli* (p < 0.0001), with mean nuclear areas of 58.70 µm^2^ and 32.22 µm^2^ respectively, compared to 35.64 µm^2^ in the no-bacteria control. DNase treatment reduced nuclear size to 27.99 µm^2^, approaching control levels, raising the possibility of altered activation states or the induction of NETosis (Fig. 3C). Representative fluorescence images show neutrophil phagocytosis of *E. coli* (Fig. 3D) and *S. aureus* (Fig. 3E) pHrodo bioparticles, with increased phagocytic activity visible under *S. aureus* conditions. Phagocytosis was quantified using pHrodo fluorescence intensity, with neutrophils showing increased uptake of *S. aureus* compared to *E. coli* (p < 0.0001) (Fig. 3F).

**Figure 3.**
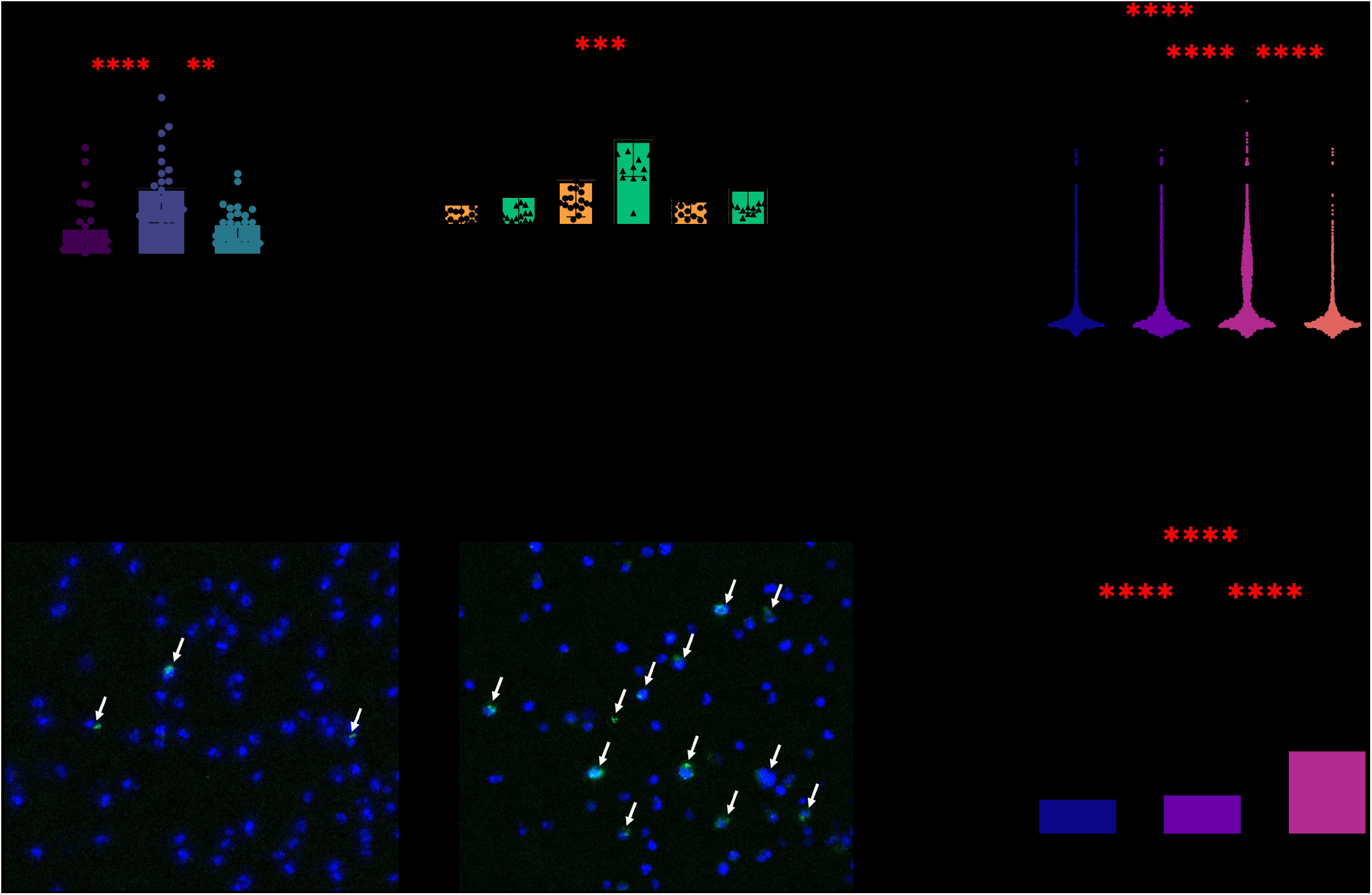
Differential neutrophil migration and phenotypic responses to *E. coli* and *S. aureus*. (A) Total neutrophil counts within the collagen matrix at 4 hours, showing significantly reduced migration in the presence of *S. aureus* compared to *E. coli*, with *S. aureus* migration indistinguishable from the no-bacteria control. (B) Proximal (P) versus distal (D) neutrophil counts relative to the lymphatic vessel at 4 hours, demonstrating preferential migration toward the vessel in the presence of *E. coli* but not *S. aureus*. (C) Neutrophil nuclear size at 24 hours, showing significantly larger nuclei in neutrophils exposed to *S. aureus* compared to *E. coli* and control conditions, a phenotype rescued by DNase treatment. (D) Representative fluorescence image of neutrophil phagocytosis of *E. coli* pHrodo bioparticles (white arrows). (E) Representative fluorescence image of neutrophil phagocytosis of *S. aureus* pHrodo bioparticles (white arrows), showing visibly increased phagocytic activity. Blue – DAPI; Green – pHrodo (bacterial uptake). (F) Quantification of pHrodo fluorescence intensity per neutrophil at 4 hours, confirming significantly greater phagocytic uptake of *S. aureus* compared to *E. coli* and control. All experiments were performed using neutrophils isolated from n = 4 independent donors. Data in A, B, and F represent mean ± SD; data in C show individual values with mean. Between-condition comparisons performed using Kruskal-Wallis with Dunn’s post-hoc test; proximal versus distal comparisons performed using Mann-Whitney test. **p < 0.01, ***p < 0.001, ****p < 0.0001; ns, not significant.

### *Staphylococcus aureus* induced NETosis

*Staphylococcus aureus*, the most common cause of wound infection, exhibited reduced neutrophil migration in our system compared to *E. coli* (Fig. 3). Given that *S. aureus* has been linked to excessive NET formation and persistence through evasion of NET-mediated killing^29–32^, we next examined whether *S. aureus* induced NETosis in our model. As described above, devices containing a biomimetic lymphatic vessel were prepared with either *E. coli* or *S. aureus* pHrodo bioparticles embedded in the collagen matrix and neutrophils were introduced into the adjacent lumen and incubated for 8 hours. NETosis was assessed by nuclear morphology using Hoechst staining and immunostaining for myeloperoxidase (MPO), a NET-associated marker. PMA, a known inducer of suicidal NETosis, served as a positive control. To evaluate the contribution of extracellular DNA, select devices were pre-treated with DNase. Neutrophils exposed to *S. aureus* displayed strong MPO positivity comparable to PMA-treated controls (Fig. 4), confirming NET-associated activation. Despite similar increases in nuclear size, distinct morphological differences were observed between conditions. Neutrophils exposed to *S. aureus* exhibited rounded or peanut-shaped nuclei, whereas PMA-treated neutrophils displayed diffuse nuclei with irregular, wispy edges characteristic of suicidal NETosis. In contrast, neutrophils exposed to *E. coli* largely retained a multilobular nuclear morphology, resembling control conditions despite the presence of pHrodo. Importantly, the enlarged nuclear phenotype induced by *S. aureus* was rescued by DNase treatment, implicating extracellular DNA as a mediator of this response.

**Figure 4.**
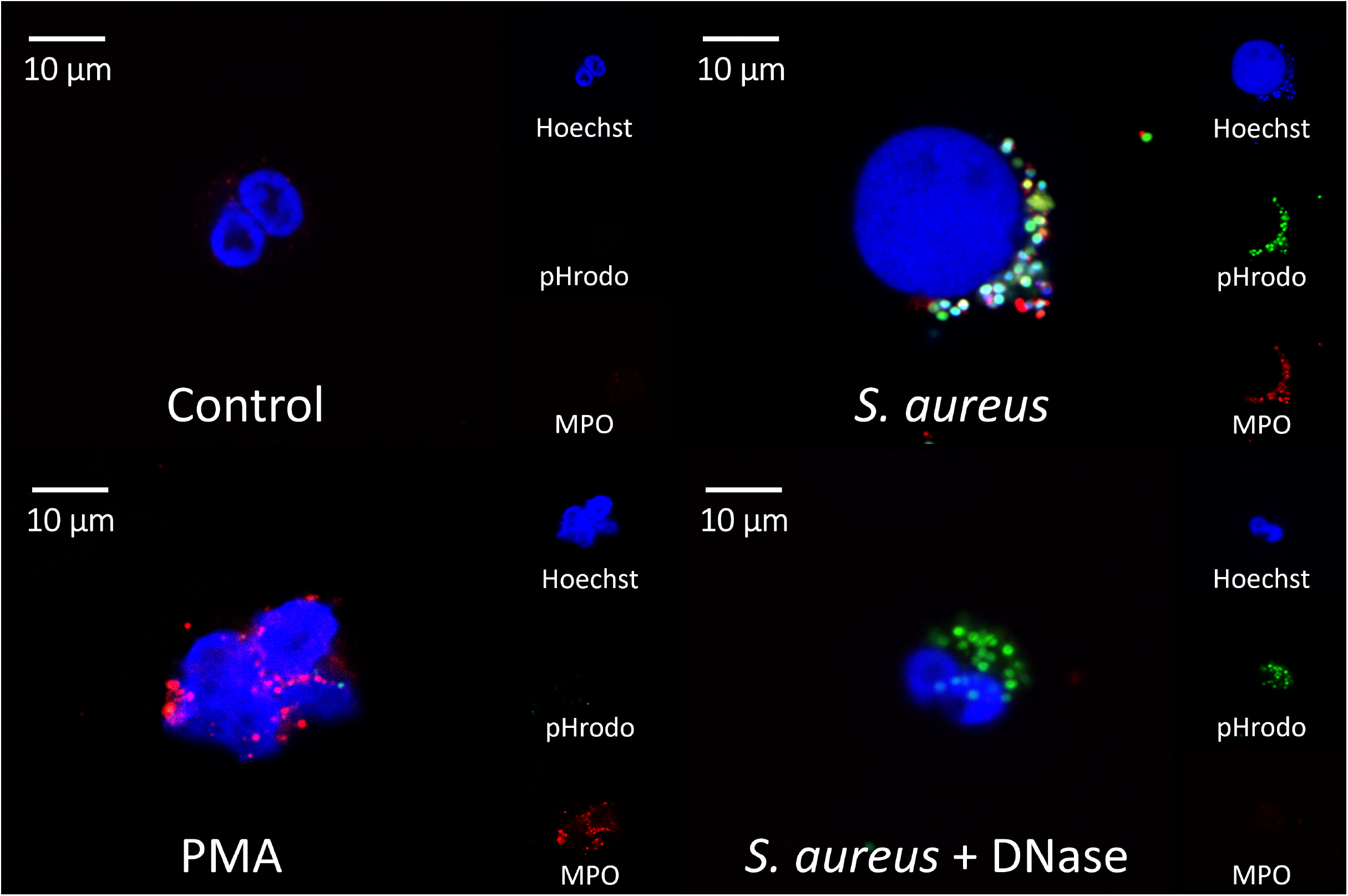
*S. aureus* induces NETosis in the lymphatic MPS. Representative confocal images (100x) of neutrophils at 8 hours under four conditions: untreated control showing intact multilobed nuclear morphology (DAPI, blue); *S. aureus* showing nuclear swelling and MPO-positive extracellular trap formation (red) alongside active phagocytosis of pHrodo bioparticles (green), consistent with vital NETosis; PMA as a positive control for suicidal NETosis showing nuclear swelling (DAPI, blue) and MPO release (red); and *S. aureus* + DNase showing rescued nuclear morphology (DAPI, blue) and reduced extracellular trap formation. Individual fluorescence channels are shown for each condition. Blue – DAPI (nuclear); Green – pHrodo (*S. aureus* bioparticles); Red – MPO.

## Discussion

Microphysiological systems (MPS) enable investigation of early immune responses in controlled, human-relevant environments that are difficult to recapitulate using conventional in vitro approaches or animal models. The LumeNEXT platform allows formation of organized luminal structures within a three-dimensional extracellular matrix and has been applied across vascular, lymphatic and cancer biology contexts^19–22^. In this study, we used this system to reconstruct a biomimetic lymphatic microenvironment and examine how lymphatic endothelium and bacterial identity influence neutrophil behavior during early inflammation. These findings indicate that lymphatic endothelial cells contribute to the local regulation of neutrophil responses during early infection, influencing recruitment dynamics and NET-associated phenotypes. By integrating luminal lymphatic endothelium with human neutrophils and bacterial stimuli, this model enabled simultaneous assessment of migration behavior, phagocytic responses, and NET-associated changes across distinct microbial conditions. This multimodal approach provides insight into how vascular context and pathogen identity together shape early inflammatory responses in human-relevant systems.

Our findings support the central hypothesis that lymphatic endothelial cells modulated neutrophil behavior in a bacterial-dependent manner. Neutrophils exhibited significantly increased migration in the presence of both bacterial stimuli and a lymphatic vessel, with a clear preference for migrating toward the lymphatic lumen. This directional migration was lost when bacterial cues were absent, underscoring the importance of infection in activating coordinated immune-lymphatic signaling. The MPS model thus recapitulates key features such as multicellular crosstalk of immune-microenvironment interactions and demonstrates the ability of HdLECs to regulate neutrophil recruitment in the context of infection. While lymphatic endothelial cells have been implicated in regulating leukocyte trafficking and facilitating neutrophil migration to lymph nodes^26,33–35^, their role in directly shaping neutrophil phenotype and function within the wound microenvironment has remained unclear. Here, we demonstrated that lymphatic endothelium amplified neutrophil recruitment and directed migration toward the vessel specifically in the presence of bacteria, revealing a previously underappreciated role for lymphatic endothelial cells in shaping innate immune responses within a 3D infection model.

We also observed that bacterial identity significantly influenced neutrophil functional phenotypes. Consistent with previous studies showing the inhibitory effects of *Staphylococcus aureus* on neutrophil migration^36^, our data revealed that *S. aureus* suppressed neutrophil migration and directionality relative to *Escherichia coli*, despite inducing a higher rate of phagocytosis. This combination of reduced migration and increased phagocytic activity raised the possibility that *S. aureus* impairs neutrophil motility following bacterial uptake, potentially limiting clearance and promoting infection persistence.

Neutrophils can respond to infection by releasing web-like chromatin structures known as neutrophil extracellular traps (NETs). *S. aureus* is known to induce excessive NET formation and possesses multiple virulence factors that allow it to evade NET-mediated killing^8,9^. Recent studies suggest that excessive NET formation is even required for *S. aureus* colonization, an effect that is rescued with DNase treatment^36^. *S. aureus* can induce both suicidal^29^ and vital NETosis^30–32^, but whether it preferentially triggers one form over the other remains unclear. *S. aureus* exposure induced a unique neutrophil phenotype characterized by increased nuclear size and positive MPO staining. The rounded or peanut-shaped nuclear morphology observed in *S. aureus*-exposed neutrophils is consistent with vital NETosis, in which nuclei remain intact with defined borders, in contrast to the diffuse, irregular nuclear dissolution characteristic of suicidal NETosis seen in PMA-treated controls. Nuclear morphology and extracellular DNA distribution are established markers used to distinguish between suicidal and vital NETosis^30–32^. The rescue of nuclear swelling following DNase treatment implicates extracellular DNA as the primary mediator and further supports the classification of this response as vital NETosis, in which DNA release, rather than cell lysis, drives the phenotype. This correlated with impaired neutrophil migration, suggesting that induction of vital NETosis by *S. aureus* may represent an immune evasion strategy that limits effective bacterial clearance.

This study was designed to isolate early neutrophil–lymphatic–microbial interactions within a controlled, human-relevant microphysiological system. As such, the platform focuses on a defined set of cell types and early inflammatory time points, enabling controlled, high-resolution analysis of neutrophil behavior. Bacterial stimuli were modeled using pHrodo-conjugated bioparticles to permit standardized assessment of phagocytosis and neutrophil activation across conditions. Nuclear morphology and MPO staining were used to assess NET-associated activation however, definitive discrimination between vital and suicidal NETosis remains technically challenging in vitro. While this approach does not capture all aspects of wound healing or long-term immune resolution, it enables direct mechanistic interrogation of acute neutrophil responses to lymphatic and microbial cues. To our knowledge, this is the first 3D microphysiological system to incorporate a biomimetic lymphatic vessel within an infected microenvironment to directly investigate neutrophil-lymphatic interactions. While existing MPS infection models have primarily utilized blood endothelial vessels to study neutrophil extravasation and migration toward bacterial stimuli^23,37^, the role of the lymphatic endothelium in shaping neutrophil responses during active infection has remained largely unexplored.

Together, these data demonstrate the utility of our 3D MPS model for dissecting complex interactions between the immune system, lymphatic vasculature and skin microbiota. The inclusion of defined microbial components—both commensal and pathogenic—enabled controlled exploration of host-microbe dynamics in early inflammation making this system suitable for a wide range of studies focused on infection, inflammation and tissue repair. This system also provides a foundation for future work exploring how microbial communities influence immune activation and lymphatic function in wound healing and supports the development of microbiome-informed strategies to improve healing outcomes and reduce antibiotic reliance. A deeper understanding of these immune–microbiome connections may ultimately guide new therapeutic approaches that enhance tissue repair while limiting dependence on systemic antibiotics.

## Methods

### Cell culture

Adult primary human dermal lymphatic endothelial cells (HdLECs) were purchased from PromoCell (C-12217). HdLECs were cultured on flasks pre-coated with 0.03mg/ml fibronectin bovine plasma (Sigma-Aldrich, F1141) at a density of 1 million cells per T75 culture flask in Endothelial Cell Growth Medium MV (ECM-MV) (PromoCell, C-22020). HdLECs were used between passage 4-6. For use in devices, cells were resuspended in Microvascular Endothelial Cell Growth Medium-2 (EGM-2MV) BulletKit (Lonza, CC-3202) at a concentration of 20,000 cells/ul. HdLECs were expanded and maintained in ECM-MV medium in accordance with manufacturer recommendations. For device fabrication, EGM-2MV was incorporated into the collagen matrix, as this medium supports collagen polymerization and allowed experimental conditions to remain consistent across all devices.

### Microphysiological device

Devices were fabricated using soft lithography, polydimethylsiloxane (PDMS) molds assembled onto glass-bottom MatTek dishes with integrated removable rods as previously described^24^. Devices possess double lumens with three devices per dish (Fig. 1). To prepare the pHrodo BioParticles, 50 µl of pHrodo™ Green BioParticles (stock concentration of 1 mg/ml) derived from *Escherichia coli* (ThermoFisher, P35366) or *Staphylococcus aureus* (ThermoFisher, P35367), depending on the experimental condition, was added to 200 µl human serum (Sigma, H6914) and 200 µl Dulbecco’s phosphate-buffered saline (DPBS) and incubated for 30 minutes at 37ºC followed by three washes with 400 µl DPBS (spun at 13,000 x g for 10 minutes). After washes, supernatant was aspirated and pHrodo BioParticles were resuspended in 300 µl EGM-2MV and added to the collagen matrix at a final concentration of approximately 0.097 mg/ml. To create the matrix in the device, a collagen mixture containing PBS, NaOH, fibronectin, EGM-2MV media (+/- pHrodo BioParticles) and collagen (9.48 mg/ml stock concentration) for a final collagen concentration of 3 mg/ml, pH 7.2-7.4 was used. Collagen was added to each device and allowed to polymerize at room temperature for 45 minutes. After polymerization, PDMS rods are removed and resultant lumens are flushed with ECM-MV media (Fig. 1A). Media is gently aspirated and fibronectin added to one lumen (designated left or right for each experiment) where the HdLECs will be placed (including controls) and incubated for 15 minutes at 37ºC. The lumen was washed twice with ECM-MV media and 2 µl of harvested HdLECs at 20,000 cells/µl were added to the fibronectin coated lumen of each device (aside from no HdLEC control) and incubated for 15-20 minutes at 37ºC before being turned upside down for a total of two rotations on each side. The HdLEC lumen was gently aspirated and flushed with fresh media then allowed to incubate overnight at 37ºC with a fresh media change in the morning, resulting in the formation of a biomimetic lymphatic vessel (Fig. 1C).

### Neutrophil isolation

Neutrophils were isolated from the blood of healthy human donors acquired under an approved Institutional Review Board protocol (University of Wisconsin-Madison, IRB #2020-1623). Cells were isolated within 1 hour of collection (blood collected in EDTA tubes) by negative selection using EasySep Direct Human Neutrophil Isolation Kit (Stemcell, 19166) following manufacturer’s protocols. Neutrophils were resuspended in ECM-MV media + 0.0002 mg/ml Hoechst to a concentration of 8,000 cells/µl for use in devices. 4 µl of the neutrophil suspension was added to the opposite lumen of each device in addition to neutrophil only controls (opposite to empty fibronectin coated lumens) for a total of 32,000 neutrophils per lumen.

### Imaging and analysis

Using an Axio Observer 7 (Zeiss) microscope, Z-stack bright field and fluorescent images were acquired to capture all neutrophils within the system. Utilizing the ImageJ image processing software, bright field and fluorescent channels were split and a z-project image was created (minimum intensity for bright field, maximum intensity for fluorescent channel). Lumens were outlined and saved as region of interests (ROIs). For fluorescent channels, the background was deleted (rollingball radius, 50) and Gaussian blur applied (SigmaValue, 1). For migration analyses, 5x Z-stack images were acquired at 20 µm increments at 4 hours post-neutrophil addition. ROIs corresponding to individual neutrophils (blue channel) were generated using StarDist (percentile range 1–99.89; probability threshold 0.4; non-maximum suppression threshold 0.4) and saved. Using the lumen ROIs, all ROIs located within either lumen were removed, leaving only neutrophils that had entered the collagen matrix. These measurements were used to quantify the number of neutrophils entering the collagen and the distance migrated. For analysis of neutrophil nuclear size and phagocytic activity, 10x Z-stack fluorescent images were acquired at 12.5 µm increments. Phagocytosis was quantified at 4 hours, while nuclear size was assessed at 24 hours following neutrophil introduction. Individual neutrophil ROIs were generated using StarDist (percentile range 1.0–99.9; probability threshold 0.4; non-maximum suppression threshold 0.4). Following lumen exclusion, nuclear area measurements were used to assess neutrophil nuclear size and pHrodo signal was quantified by overlaying neutrophil ROIs onto the green channel and measuring fluorescence intensity. All measurements were plotted using GraphPad Prism. Statistical analyses were performed using Kruskal-Wallis tests to compare conditions and Mann-Whitney tests for within-condition comparisons.

### MPO Staining and NETosis analysis

Devices were prepared as previously described, with a biomimetic lymphatic vessel formed by human dermal lymphatic endothelial cells (HdLECs) in one lumen and isolated human neutrophils (stained with Hoechst) introduced into the opposing lumen. The collagen matrix contained either *Escherichia coli* or *Staphylococcus aureus* pHrodo-conjugated bioparticles as a bacterial stimulus at a concentration of approximately 0.097 mg/ml, or no-bacteria as a control. A suicidal NETosis positive control was included using phorbol 12-myristate 13-acetate (PMA, Abcam). A 100 µM PMA stock solution was serially diluted to a final working concentration of 1 µM. For device treatment, 5 µL of 1 µM PMA was added to 495 µL of EGM-MV media, added to both lumens and incubated for 15 minutes at 37 °C. In parallel, neutrophils were pre-incubated with PMA by adding 2.5 µL of 1 µM PMA to 247.5 µL of neutrophils (8,000 cells/µL) for 15 minutes at 37 °C prior to loading into the empty lumen. To assess the contribution of extracellular DNA in NETosis, an additional DNase condition was included. DNase I (STEMCELL Technologies, 07900) was prepared by diluting 5 µL of a 1 mg/mL stock into 95 µL of EGM-MV medium for device treatment. Separately, neutrophils (95 µL at 8,000 cells/µL) were treated by adding 5 µL of DNase I stock directly to the cell suspension. Both DNase-treated devices and neutrophils were incubated for 15 minutes at 37 °C prior to neutrophil loading. At 8 hours post-neutrophil introduction, devices were fixed and stained. Lumens were washed twice with DPBS, fixed with 4% paraformaldehyde for 15 minutes and permeabilized with 0.2% Triton X-100 for 30 minutes at room temperature. Following three 0.1% DPBS + Tween (DPBST) washes, devices were flooded with blocking buffer containing 3% BSA, 5% human serum, 0.1 M glycine in 0.1% PBST, then incubated overnight at 4 °C. Devices were incubated overnight at 4 °C with myeloperoxidase (MPO) recombinant rabbit monoclonal antibody (1 mg/mL, MA5-32582) diluted 1:100 in staining buffer (3% BSA in 0.1% DPBST). The following day, devices were washed five times for 10 minutes each with 0.1% DPBST, flooded with 0.1% DPBST and stored overnight at 4 °C. Secondary antibody incubation was performed using goat anti-rabbit Alexa Fluor 568 (A1106) diluted 1:100 in 0.1% DPBST containing 10% goat serum for 1 hour at room temperature. Devices were washed 3-4 times for 20 minutes each, followed by an overnight wash at 4 °C prior to imaging. Imaging was performed at 100x magnification using a Nikon Yokogawa W1 spinning disk confocal microscope.

## Acknowledgements

This work was supported by the National Institutes of Health Training Grant 5T32OD010423. We thank the Optical Imaging Core at the University of Wisconsin–Madison for use of the Nikon Spinning Disk Confocal Microscope. This research was also supported in part by the University of Wisconsin Carbone Cancer Center Support Grant P30CA014520. This work was also supported by NIH R01AI154940 and R01AI172874.

## Author contributions statement

K.A.S. conceived the study, designed and conducted the experiments, analyzed the results, and wrote the manuscript. S.C.K. supervised the work and contributed to manuscript editing. D.J.B. supervised the work, provided resources, and contributed to manuscript editing. All authors reviewed the manuscript.

## Additional information

David J. Beebe holds equity in Bellbrook Labs LLC, Salus Discovery LLC, Lynx Biosciences Inc., Stacks to the Future LLC, Flambeau Diagnostics LLC, Eolas Diagnostics, Inc., Navitro Biosciences LLC, and Onexio Biosystems LLC.

## Data Availability

All data supporting the findings of this study are available within the paper and its Supplementary Information. Additional datasets generated during the current study are available from the corresponding author upon reasonable request.

## References

1. Mainous III, A. G., Diaz, V. A., Matheson, E. M., Gregorie, S. H. & Hueston, W. J. Trends in Hospitalizations with Antibiotic-Resistant Infections: U.S., 1997–2006. Public Health Rep 126, 354–360 (2011).

2. Baggs, J. et al. Risk of Subsequent Sepsis Within 90 Days After a Hospital Stay by Type of Antibiotic Exposure. Clin Infect Dis 66, 1004–1012 (2018).

3. Leliefeld, P. H., Wessels, C. M., Leenen, L. P., Koenderman, L. & Pillay, J. The role of neutrophils in immune dysfunction during severe inflammation. Crit Care 20, (2016).

4. Dobrewa, W., Bielska, M., Bąbol-Pokora, K., Janczar, S. & Młynarski, W. Congenital neutropenia: From lab bench to clinic bedside and back. Mutat Res Rev Mutat Res 793, (2024).

5. Danne, C., Skerniskyte, J., Marteyn, B. & Sokol, H. Neutrophils: from IBD to the gut microbiota. Nat Rev Gastroenterol Hepatol 21, 184–197 (2024).

6. Fresneda Alarcon, M., McLaren, Z. & Wright, H. L. Neutrophils in the Pathogenesis of Rheumatoid Arthritis and Systemic Lupus Erythematosus: Same Foe Different M.O. Front Immunol 12, 649693 (2021).

7. Klevens, R. M. et al. Estimating Health Care-Associated Infections and Deaths in U.S. Hospitals, 2002. Public Health Rep 122, 160–166 (2007).

8. Menegazzi, R., Decleva, E. & Dri, P. Killing by neutrophil extracellular traps: fact or folklore? Blood 119, 1214– 1216 (2012).

9. Berends, E. T. et al. Nuclease expression by Staphylococcus aureus facilitates escape from neutrophil extracellular traps. J Innate Immun 2, 576–586 (2010).

10. von Köckritz-Blickwede, M. & Winstel, V. Molecular Prerequisites for Neutrophil Extracellular Trap Formation and Evasion Mechanisms of Staphylococcus aureus. Front Immunol 13, 836278 (2022).

11. Speziale, P. & Pietrocola, G. Staphylococcus aureus induces neutrophil extracellular traps (NETs) and neutralizes their bactericidal potential. Comput Struct Biotechnol J 19, 3451–3457 (2021).

12. Humphreys, H., Becker, K. & Dohmen, P. M. Staphylococcus aureus and surgical site infections: benefits of screening and decolonization before surgery. J Hosp Infect 94, 295–304 (2016).

13. Khosravi, A., Yáñez, A. & Price, J. G. Gut microbiota promote hematopoiesis to control bacterial infection. Cell Host Microbe 15, 374–381 (2014).

14. Kimura, T., Sugaya, M., Blauvelt, A., Okochi, H. & Sato, S. Delayed wound healing due to increased interleukin-10 expression in mice with lymphatic dysfunction. J Leukoc Biol 94, 137–145 (2013).

15. Zhang, D. & Frenette, P. S. Cross talk between neutrophils and the microbiota. Blood 133, 2168–2177 (2019).

16. Alexeev, E. E., Dowdell, A. S. & Henen, M. A. Microbial-derived indoles inhibit neutrophil myeloperoxidase to diminish bystander tissue damage. FASEB J 35, (2021).

17. Wang, H., Chan, M. W. M., Chan, H. H. & Pang, H. Longitudinal Changes in Skin Microbiome Associated with Change in Skin Status in Patients with Psoriasis. Acta Derm Venereol 100, (2020).

18. Leystra, A. A. & Clapper, M. L. Gut Microbiota Influences Experimental Outcomes in Mouse Models of Colorectal Cancer. Genes (Basel) 10, 900 (2019).

19. Liang, N. W., Wilson, C. & Davis, B. Modeling lung endothelial dysfunction in sepsis-associated ARDS using a microphysiological system. Physiol Rep 12, (2024).

20. Gong, M. M. et al. Human organotypic lymphatic vessel model elucidates microenvironment-dependent signaling and barrier function. Biomaterials 214, (2019).

21. van Dijk, C. G. M. et al. A new microfluidic model that allows monitoring of complex vascular structures and cell interactions in a 3D biological matrix. Lab Chip 20, 1827–1844 (2020).

22. Virumbrales-Muñoz, M. et al. Organotypic primary blood vessel models of clear cell renal cell carcinoma for single-patient clinical trials. Lab Chip 20, 4420–4432 (2020).

23. Hind, L. E., Ingram, P. N., Beebe, D. J. & Huttenlocher, A. Interaction with an endothelial lumen increases neutrophil lifetime and motility in response to P aeruginosa. Blood 132, 1818–1828 (2018).

24. Jiménez-Torres, J. A., Peery, S. L., Sung, K. E. & Beebe, D. J. LumeNEXT: A Practical Method to Pattern Luminal Structures in ECM Gels. Adv Healthc Mater 5, 198–204 (2016).

25. Yang, Q., Langston, J. C. & Prosniak, R. Distinct functional neutrophil phenotypes in sepsis patients correlate with disease severity. Front Immunol 15, (2024).

26. Özcan, A., Collado-Diaz, V. & Egholm, C. CCR7-guided neutrophil redirection to skin-draining lymph nodes regulates cutaneous inflammation and infection. Sci Immunol 7, (2022).

27. Xue, J., Lin, Y. & Oo, D. Lymph-derived chemokines direct early neutrophil infiltration in the lymph nodes upon Staphylococcus aureus skin infection. Proc Natl Acad Sci U S A 119, (2022).

28. Arokiasamy, S. et al. Endogenous TNFα orchestrates the trafficking of neutrophils into and within lymphatic vessels during acute inflammation. Sci Rep 7, (2017).

29. Wise, A. D., TenBarge, E. G. & Mendonça, J. D. C. Mitochondria sense bacterial lactate and drive release of neutrophil extracellular traps. Cell Host Microbe 33, 341–357 (2025).

30. Malachowa, N., Kobayashi, S. D., Freedman, B., Dorward, D. W. & DeLeo, F. R. Staphylococcus aureus leukotoxin GH promotes formation of neutrophil extracellular traps. J Immunol 191, 6022–6029 (2013).

31. Pilsczek, F. H., Salina, D. & Poon, K. K. A novel mechanism of rapid nuclear neutrophil extracellular trap formation in response to Staphylococcus aureus. J Immunol 185, 7413–7425 (2010).

32. Mazzoleni, V., Zimmermann, K., Smirnova, A., Tarassov, I. & Prévost, G. Staphylococcus aureus Panton-Valentine Leukocidin triggers an alternative NETosis process targeting mitochondria. FASEB J 35, (2021).

33. Hampton, H. R., Bailey, J., Tomura, M., Brink, R. & Chtanova, T. Microbe-dependent lymphatic migration of neutrophils modulates lymphocyte proliferation in lymph nodes. Nat Commun 6, (2015).

34. Gorlino, C. V., Ranocchia, R. P. & Harman, M. F. Neutrophils exhibit differential requirements for homing molecules in their lymphatic and blood trafficking into draining lymph nodes. J Immunol 193, 1966–1974 (2014).

35. Rigby, D. A., Ferguson, D. J., Johnson, L. A. & Jackson, D. G. Neutrophils rapidly transit inflamed lymphatic vessel endothelium via integrin-dependent proteolysis and lipoxin-induced junctional retraction. J Leukoc Biol 98, 897–912 (2015).

36. Bitschar, K., Staudenmaier, L. & Klink, L. Staphylococcus aureus Skin Colonization Is Enhanced by the Interaction of Neutrophil Extracellular Traps with Keratinocytes. J Invest Dermatol 140, 1054–1065 (2020).

37. Kim, J. J. et al. A microscale, full-thickness, human skin on a chip assay simulating neutrophil responses to skin infection and antibiotic treatments. Lab Chip 19, 3094–3103 (2019).

