## Supplemental Figure 1 for "Lymphatic Endothelial Cells Regulate Neutrophil Phenotypes and Function in a Microphysiological Model of Infection"

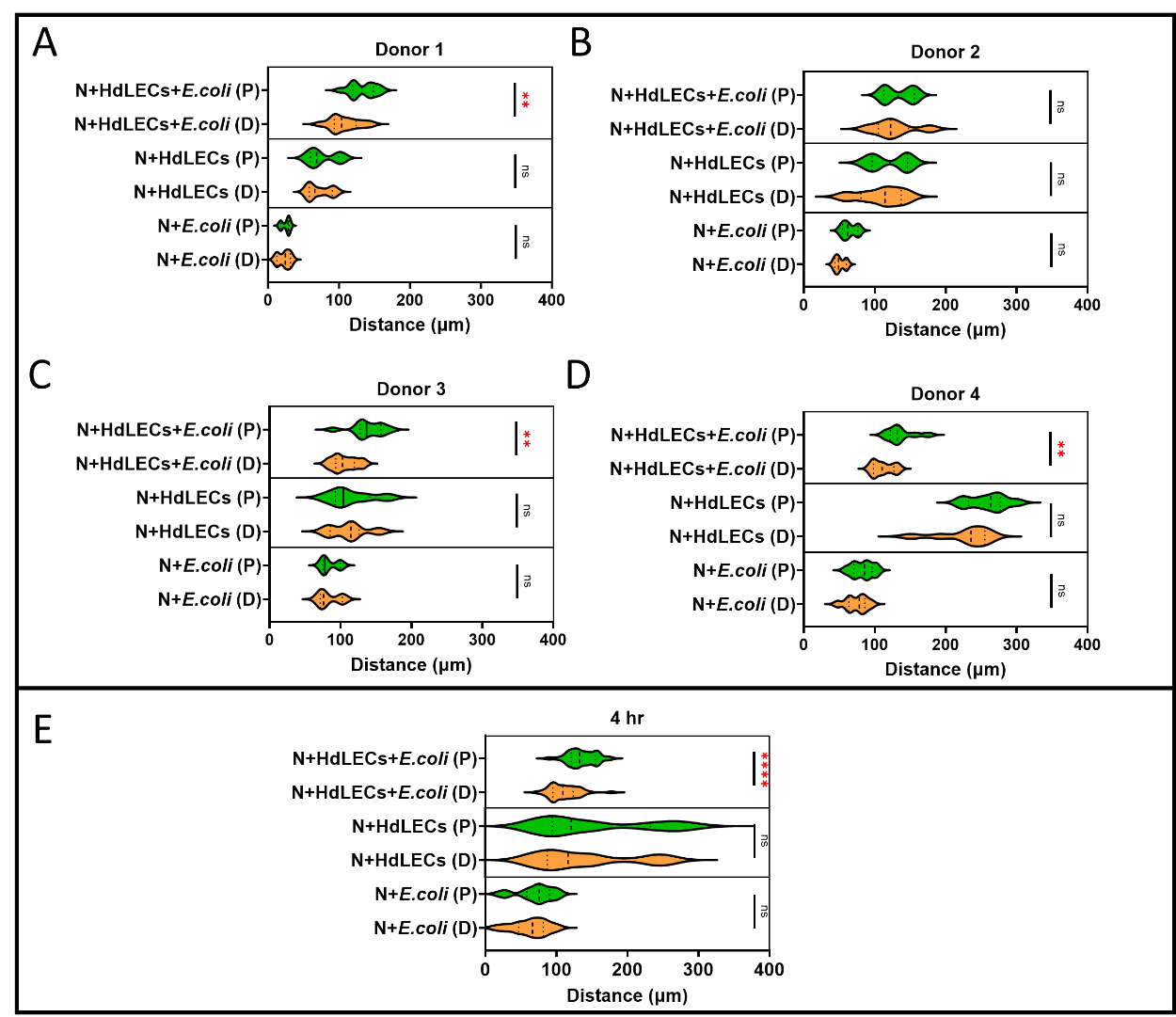

**Supplemental Figure 1. Donor-dependent neutrophil migration toward *E. coli* in the presence of lymphatic endothelial cells.** Proximal (P) versus distal (D) migration distance in the presence of *E. coli* pHrodo bioparticles, lymphatic vessel, or both, shown for individual donors (A–D) and combined (E). Preferential proximal migration in the N+HdLECs+*E. coli* condition was significant in 3 of 4 donors and in the pooled analysis. Data are displayed as violin plots. Proximal versus distal comparisons performed using Mann-Whitney test. **p < 0.01, ****p < 0.0001; ns, not significant.
